# Dorsal-to-ventral imbalance in the superior longitudinal fasciculus mediates methylphenidate’s effect on beta oscillations in ADHD

**DOI:** 10.1101/2020.09.23.309526

**Authors:** C. Mazzetti, C. G. Damatac, E. Sprooten, N. ter Huurne, J.K. Buitelaar, O. Jensen

**Affiliations:** Radboud University Nijmegen, Donders Institute for Brain, Cognition and Behaviour, Nijmegen, The Netherlands; Centre for Human Brain Health, School of Psychology, University of Birmingham, Birmingham, UK; Department of Cognitive Neuroscience, Radboudumc, Nijmegen, The Netherlands; Karakter Child and Adolescent Psychiatry University Centre, Nijmegen, The Netherlands

**Keywords:** Brain oscillations, MEG, ADHD, DTI, Biomarker, Methylphenidate

## Abstract

**Background:** While pharmacological treatment with Methylphenidate (MPH) is a first line intervention for ADHD, its mechanisms of action have yet to be elucidated. In a previous MEG study, we demonstrated that MPH in ADHD normalizes beta depression in preparation to motor responses (1). We here seek to identify the white matter tracts that mediate MPH’s effect on beta oscillations.

**Methods:** We implemented a double-blind placebo-controlled crossover design, where boys diagnosed with ADHD underwent behavioral and MEG measurements during a spatial attention task while on and off MPH. Results were compared with an age/IQ-matched typically developing (TD) group performing the same task. Estimates of white matter tracts were obtained through diffusion tensor imaging (DTI). Based on aprioristic selection model criteria, we sought to determine the fiber tracts associated with electrophysiological, behavioral and clinical features of attentional functions.

**Results:** We identified three main tracts: the anterior thalamic radiation (ATR), the Superior Longitudinal Fasciculus (‘parietal endings’) (SLFp) and Superior Longitudinal Fasciculus (‘temporal endings’) (SLFt). ADHD symptoms severity was associated with lower fractional anisotropy (FA) within the ATR. In addition, individuals with relatively higher FA in SLFp compared to SLFt showed faster and more accurate behavioral responses to MPH. Furthermore, the same parieto-temporal FA gradient explained the effects of MPH on beta modulation: subjects with ADHD exhibiting higher FA in SLFp compared to SLFt also displayed greater effects of MPH on beta power during response preparation.

**Conclusions:** Based on MPH’s modulatory effects on striatal dopamine levels, our data suggest that the behavioral deficits and aberrant oscillatory modulations observed in ADHD depend on a structural connectivity imbalance within the SLF, caused by a diffusivity gradient in favor of temporal rather than parietal, fiber tracts.

## Introduction

The neural mechanisms underlying selective attention processes are contingent upon a complex interaction of brain networks. Prior studies using electro- and magneto-encephalography (EEG/MEG) have provided insights into diverse oscillatory patterns indexing distractor suppression and appropriate target processing. These start prior to the display of the stimuli, when participants prepare to the response by specifically allocating attentional resources in order to meet task demands (2–5). One important function is the preparation to the motor response to a cued target (e.g., button press), which interacts with visual attentional allocation. The electrophysiological correlate underlying this behavioral component is visible as a suppression of oscillations in sensorimotor areas. Specifically, modulation of brain activity in the beta band (15 – 30Hz) has been associated with top-down control of motor preparation, reflecting an interaction between cognitive and motor functions, which has been interpreted as increased excitability of task-relevant brain areas (6,7). This motivates the investigation of brain oscillations during attention tasks in populations with attentional problems.

Attention deficit-hyperactivity disorder (ADHD) is a neurodevelopmental disorder characterized by age-inappropriate levels of inattention and hyperactivity-impulsivity (American Psychiatric Association, 2013). Aberrant modulation of oscillatory activity has indeed been observed in individuals identified with ADHD, both in adults (9–11) and pediatric populations (12–14), with the latter representing the most prominent line of research, given the predominance of the disorder in early life (15,16). Weaker suppression of β-band activity (13 – 30Hz) has been found in ADHD, prior to response preparation to a cued target (17), reflecting a lack of appropriate motor planning. Furthermore, recent evidence has observed weakened beta modulation during working memory encoding in ADHD (18) consistent with the view that modulation of beta oscillations reflect cognitive functions (19,20).

Previously, we showed that beta depression in preparation to responses to a cued target is reduced in children with ADHD as compared to matched TD peers (1). Additionally, in a double-blind randomized placebo-controlled design within the ADHD group we showed that MPH restores levels of beta depression in children with ADHD, by normalizing its values to the ones observed in the TD group. This effect is allegedly attributable to the effects of psychostimulants on catecholamines’ levels in the midbrain (21). Specifically, MPH blocks the reuptake of norepinephrine and, particularly, dopamine at the synaptic cleft (22–24) increasing their availability. A dopaminergic imbalance within networks mediated by the prefrontal cortex has indeed been proposed to underlie symptoms of attentional deficits and hyperactivity (25,26). Consistently, the modulatory action of dopamine is alleged to mediate the interaction between frontoparietal and default mode attentional networks (27).

So far, the link between beta oscillations and brain structure remains yet to be elucidated. This can be achieved by means of diffusion tensor imaging (DTI), which estimates the direction of diffusion of water molecules in the brain and is thought to reflect the underlying microstructural properties of white matter fiber tracts (28,29). Among the DTI-derived metrics is fractional anisotropy (FA), reflecting coherence of diffusion of water along the main tract direction (30,31) and, allegedly, the underlying tissue microstructure, such as integrity of myelin sheath, which impacts the overall mobility of water along axons (32). DTI studies in ADHD have so far pointed to reduced white matter integrity in widespread areas in the brain (33–35). Diffusion in the angular bundle of the cingulum correlates with hyperactivity-impulsivity symptomatology and associations of FA with ADHD are not uniformly distributed across white matter tracts (36). Importantly, the Superior Longitudinal Fasciculus, a white matter tract known to sustain spatial attention functions (38,39), has been consistently reported among the structural correlates of ADHD symptoms. In particular, lower integrity (FA) along this bundle has been associated with severity of ADHD symptoms and behavioral performance on cognitive tests (40–42). Taking advantage of the spatial resolution offered by MEG and the link to microstructural connectivity properties offered by DTI, this study aims to study the association between oscillatory and structural features in relation to attentional impairments in children with ADHD and typically developing (TD) children. We hence coupled the previously reported electrophysiological results (1) with an analysis of white matter microstructural properties in the same subjects.

## Methods and Materials

A detailed explanation of participants’ inclusion and exclusion criteria, attentional task, and study design has been previously described (1) and can be found in *Supplementary Materials*.

### Participants

The study included 27 children diagnosed with ADHD and 27 typically developing (TD) male children. A total of 10 children (9 ADHD) withdrew from the experiment after at least one session, due to unwillingness to proceed and/or excessive complications during testing. As a result, structural and diffusion-weighted MRI scans and MEG data were acquired for a total of 49 (26 TD) and 46 participants (26 TD), respectively.

### Experimental design

The study design is outlined in **Figure 1**. Overall, children in the ADHD group visited the lab three times, while children in the TD group twice. During the first intake-session, participants from groups underwent behavioral and IQ testing. For the ADHD group, a further in-depth intake was conducted to determine the medication dosage to be used during the task, based on operating procedures followed in prior studies (43). Based on the screening, a standardized dosage was chosen (either 10 or 15mg Methylphenidate immediate release; IR-MPH). Following the behavioral screenings, the MRI session took place for a duration of ~30mins.

**Figure 1.**
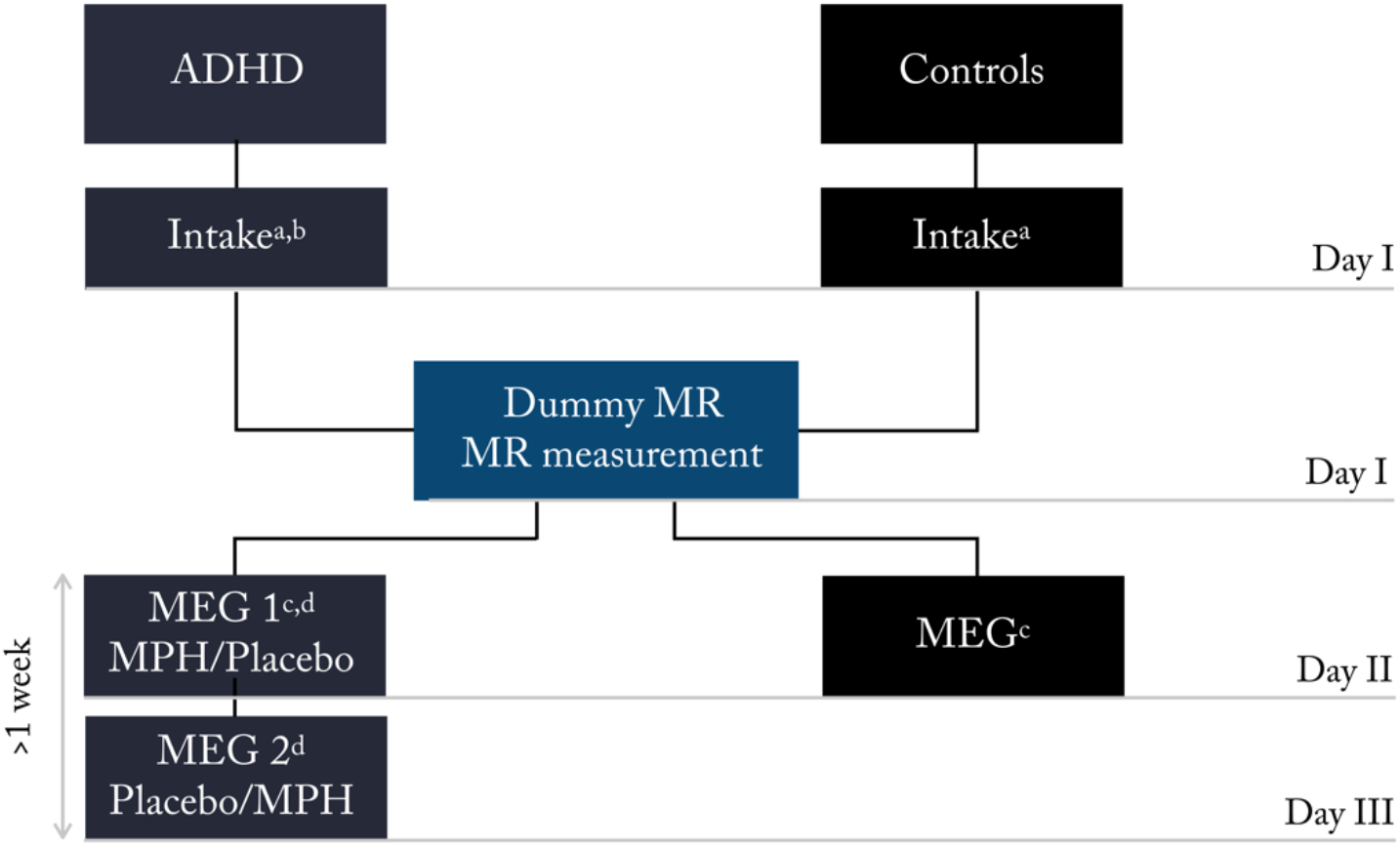
Experimental study design. ^a^ Case Report Form and Behavioural test (line bisection task, WISCIII *Vocabulary* and *Block design* subscales, ADHD rating scale, CBCL); ^b^ Psychiatric intake (basic medical screening and dosage determination); ^c^ Polhemus digitizer; ^d^ Medication intake (1 hour prior to the beginning of experimental task); Participants in the ADHD and Controls group visited the laboratory three and two times, respectively. During the first visit (day I) the psychiatric assessment took place, which determined participants’ suitability for the study. The same day, the dummy MR took place, followed by the actual MR scan. The second day (day II) both groups performed the attentional task while electromagnetic activity was recorded with the MEG. The ADHD group performed the task twice (day II and day III), once upon administration of MPH, and once upon administration of a placebo pill. This was done according to a randomized crossover design, half of the participants received the MPH on day II,

During the second visit, both groups undertook the MEG testing. For the TD group, this constituted the last day of testing, while for the ADHD group, two MEG sessions were planned at two different visits, separated by at least one-week interval. The ADHD group performed the MEG task twice, i.e., under two conditions (MPH and placebo), according to a double-blind placebo controlled randomized design. Prior to each MEG session, a 24 hour treatment suspension allowed to control for withdrawal symptoms related to drug administration (rebound effect) (44). MEG testing began one hour after medication intake, allowing to reach on average moderate plasma concentration (*C*_max_) of the drug along the experiment, which progressively increases and reaches its peak around the second hour post-intake (45). After completion of the MEG session, participants proceeded with their own regular stimulant.

### The attention task

The task was presented as a child-friendly adaptation of a Posner’s cueing paradigm for spatial orienting of attention, where a central cue (represented with a clown fish looking either at the left or at the right side of the screen) indicated the upcoming position of a target (a shark with an open mouth). Participants were asked to indicated via button press the position of the target, while ignoring the distractor on the opposite screen side (shark with mouth closed) (see **Figure 2)**.

**Figure 2.**
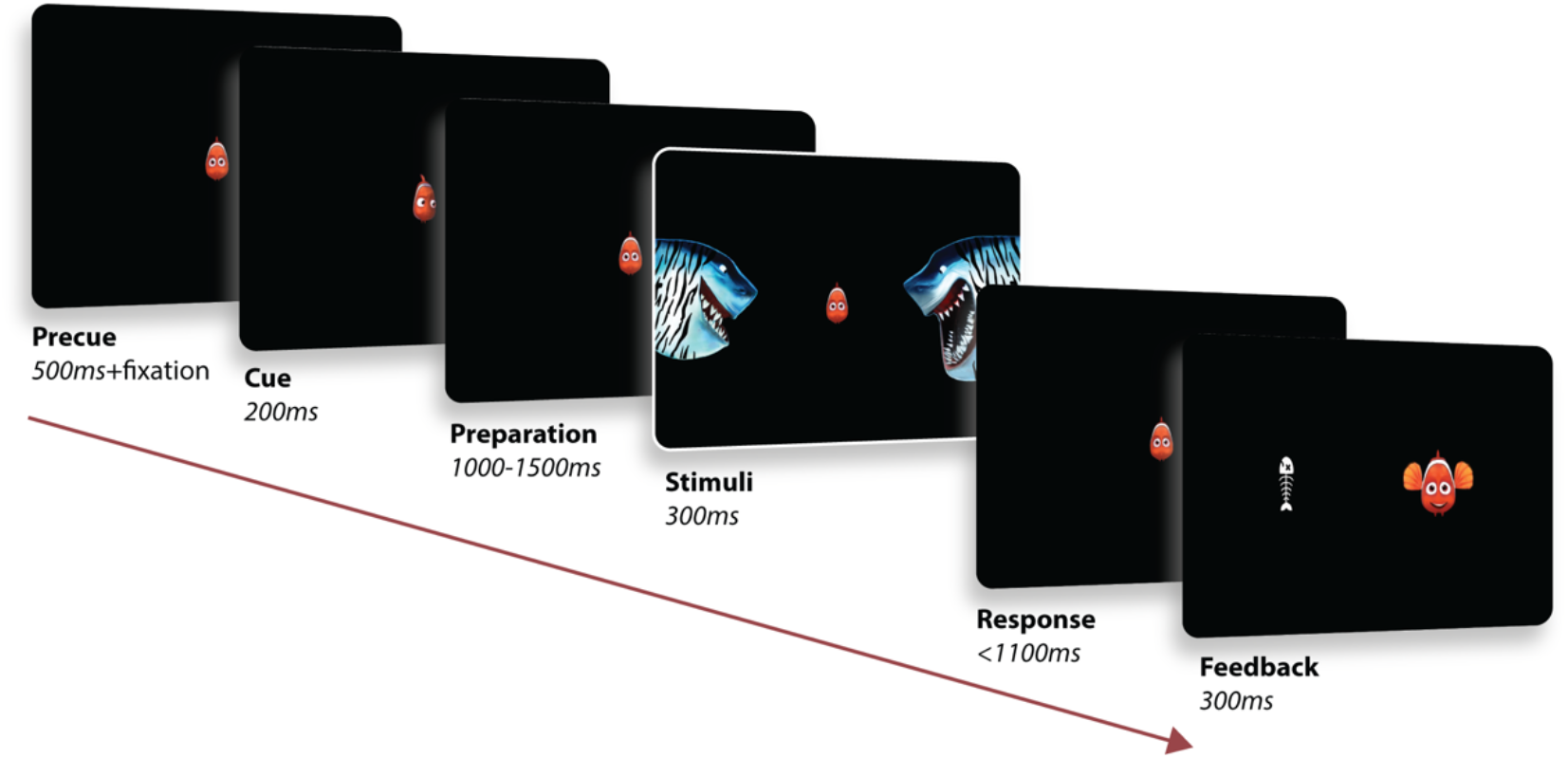
Attentional task. (Adapted from Mazzetti et al., 2020). The paradigm consisted in a child-friendly adaptation of a Posner cueing paradigm for the study of spatial orienting of attention. Each trial (370 in total) began with the presentation of a fish in the middle of the screen, serving as fixation cross. An eye tracker ensured that the children kept proper fixation throughout the whole trial (as trials were stopped in case the subject performed a saccade). A cue was then presented for 200ms, represented by the fish looking either at the left or right side of the screen (cue side equally distributed across trials). After a preparation interval jittered in the range 1100 – 1500ms, the stimuli were then presented for 300ms, on the two sides of the screen. The child was asked to respond, via button press, indicating the position of the target (shark with an open mouth), while ignoring the distractor on the other side (shark with mouth closed). A positive feedback (happy fish) was then presented if a correct answer was provided within the response interval (1100ms). In case of wrong or no response, a negative feedback was presented (fish bone).

The experiment consisted of 360 trials, equally divided in 10 blocks, after which the participant was given the possibility to take a break and/or talk to the parents. Note that, for the ADHD group, the treatment order for the two MEG sessions was randomized across participants.

### MEG data acquisition and analysis

Electromagnetic brain activity was recorded from the participants seated in a CTF 275-sensor whole-head MEG system with axial gradiometers (CTF MEG Systems, VSM MedTech Ltd.). Head position was monitored throughout the experiment via online head-localization software, allowing, if necessary, readjustment of the participant’s position between blocks. Complete illustration of the steps followed for the analysis of power and beta oscillatory indices of interest, can be found in our previous paper (1) and are further described in the Supplementary Materials.

Importantly, in the above study, we report how the preparation to motor response to the cued target was accompanied by a desynchronization of beta oscillations in the MEG data at central sensors. For each subject, a Beta Preparation Index (PI(β)) was hence computed by considering the average beta band modulation over the sensors and time window of interest (f= 15 – 30Hz, −1000 < t < 0 ms).

To estimate MPH modulatory effects exerted on beta preparation, we considered the difference in PI(β)) between MPH drug and placebo in the ADHD sample, referred to as ΔPI(β):

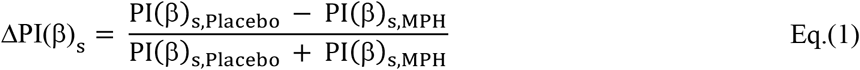

Based on Eq.1, the larger the ΔPI(β) the higher the stronger in the beta depression following MPH intake.

### Structural data analysis

MRI and DTI data were acquired via a 3T MAGNETOM Skyra MR scanner (Siemens AG, Healthcare Sector, Erlangen, Germany) with a product 32-channel head coil. The protocol included a T1-weighted MRI scan for anatomical reference and analysis and diffusion-weighted MRI scans for performing fiber tractography.

Anatomical and DTI images were analyzed in FreeSurfer 6.0 (http://surfer.nmr.mgh.harvard.edu/). The TRACULA toolbox (Tracts Constrained by Underlying Anatomy; Yendiki et al., 2011) was implemented for preprocessing of DWI images and for subsequent delineation of 18 major white matter tracts (8 bilateral and 2 interhemispheric): corticospinal tract (CST), inferior longitudinal fasciculus (ILF), uncinate fasciculus (UNC), anterior thalamic radiations (ATR), cingulum-cingulate gyrus bundle (CCG), cingulum-angular bundle (CAB), superior longitudinal fasciculus-parietal terminations (SLFP), superior longitudinal fasciculus-temporal terminations (SLFT), corpus callosum forceps major and minor (Fmaj, Fmin). TRACULA allows the automated reconstruction of major white matter pathways based on the global probabilistic approach described in (47), and further extends it by incorporating anatomical knowledge in the prior probability function: each resulting segmented tract is the best fit not only given the observed diffusion data within each subject, but also given its similarity to the known tract anatomy in relation to grey matter segmentations from FreeSurfer.

Preprocessing steps included eddy current compensation, head motion correction, intra-subject registration (individual DWI to individual T1), inter-subject registration (individual T1 affine registration to the MNI template space), creation of cortical masks (parcellation via probabilistic information estimated from a manually labeled training set based on Desikan-Killiany Atlas, **Figure 3**) and white-matter masks (based on (44)), tensor fitting for extraction of tensor-based measures, computation of anatomical priors for white-matter pathways reconstruction (e.g., diffusion eigenvectors, eigenvalues and FA for each voxel). Next, *bedpostx* was applied with a two-fiber, ball-and-stick model to estimate the distributions of the diffusion parameters and create the input for probabilistic tractography (49).

**Figure 3.**
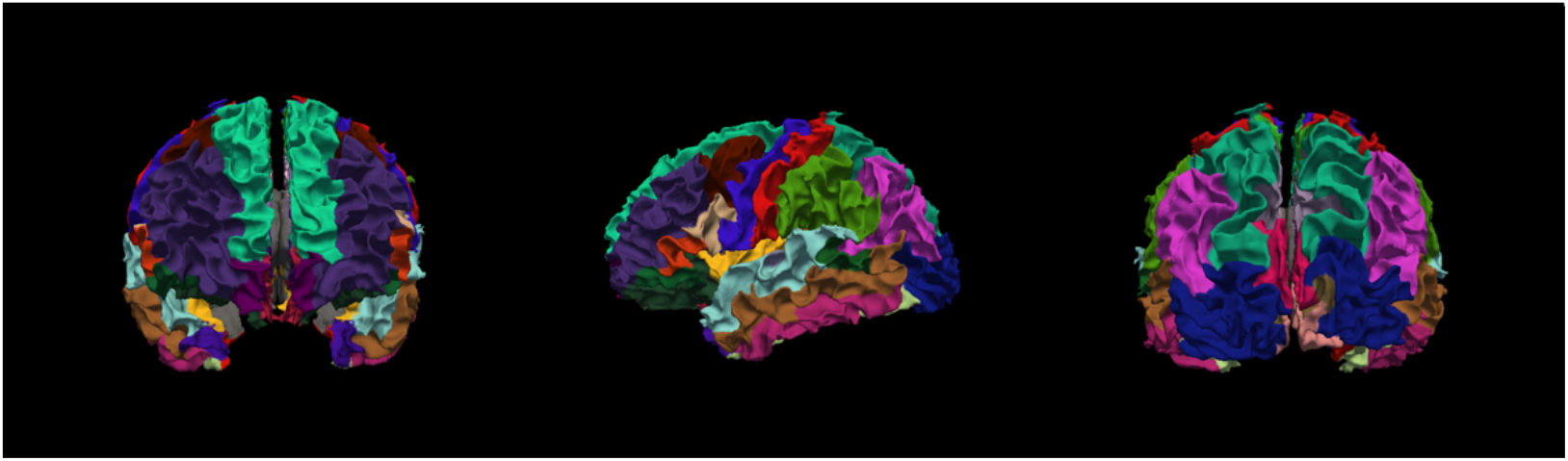
3D rendering of cortical parcellation based on Desikan-Killiany atlas in one sample TD subject. A cortical parcellation was generated prior to tract estimation, which are combined with prior distributions on the neighboring anatomical structures of each pathway and subcortical segmentation to constrain the tractography solutions (obviating the need for user interaction thus automating the process).

See **Figure 4** for an example of TRACULA’s output in a single healthy TD participant, showing the posterior distribution for all the white matter pathways included in the segmentation pipeline. Each participant’s scan was then registered to standard MNI space for group-level analyses. A pairwise correlation matrix for bilateral structures is presented in **Figure 5**, showing that no negative association was present between tracts’ FA values.

**Figure 4.**
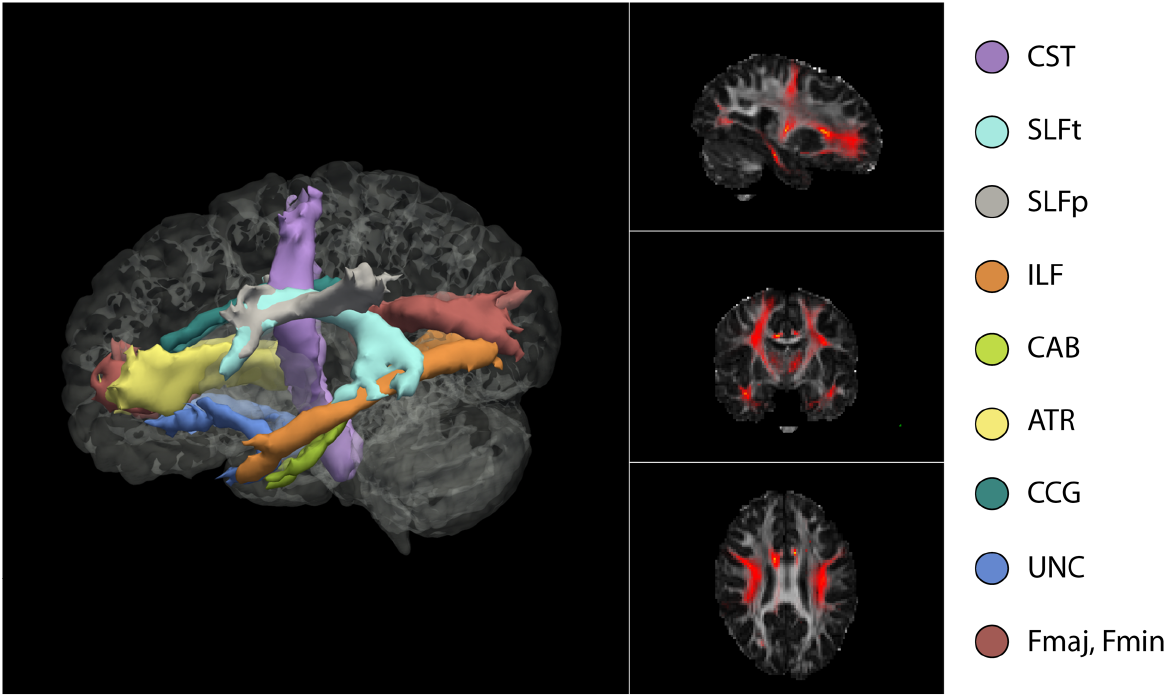
3D isosurface rendering and 2D orthographic view of tract reconstructions in one sample subject obtained using TRACULA. Visualization of the probability distributions of all whitematter pathways simultaneously overlaid on 4D brain mask. All 18 tracts are displayed at 20% of their maximum threshold

**Figure 5.**
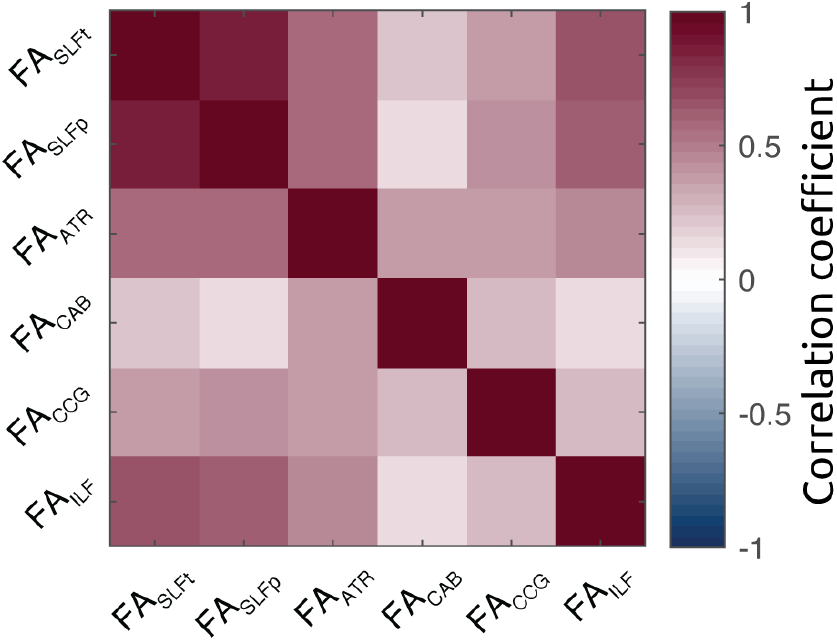
Correlation matrix of FA across bilateral white matter structures shows that no negative association subsists between segmented tracts.

All statistical analyses were performed in MATLAB2019a. For all analyses, the main metric of interest was mean FA within the tracts.

#### Model selection of white matter ROIs

In order to determine the relationship between mean FA along the white matter tracts segmented, as well as electrophysiological and behavioural measures, we implemented a general linear model (GLM) specifying the FA values of the white matter structures as regressors for the prediction of each index of interest.

Consistent with the methods implemented in previous work (50), and given the relative heterogeneity of current results in the field (33,51,52), we implemented a data driven strategy aimed at selecting the optimal set of or regressors to be included in the model.

We started by focusing on the behavioral performance to constrain the optimal model, hence setting IESs as dependent variable. Next, we considered all possible combinations from 2 to 5 regressors reflecting the FA values of the bilateral tracts segmented with TRACULA: SLFt, SLFp, ATR, CAB, CCG, ILF (in addition to the model including all 6 regressors, referred to as the ‘full model’). We hence separately considered the general linear mixed models (maximum likelihood estimation) derived from all possible combinations of *n* regressors, i.e., including either 2, 3, 4 or 5 regressors (i.e. ROIs), by computing all possible unique permutations of *n* regressors from the subset defined. This resulted in a set of models for each of the 4 possibilities using 2, 3, 4 or 5 regressors. Next, for each of these subsets, we derived the model associated with the lowest *Akaike Information Criterion* (AIC), *Bayesian Information Criterion* (BIC), highest *log likelihood* (where the winning model would be the one associated with at least two highest criteria compared to the others in the same model subset). These values have been commonly used in model selection to identify the best predictor subsets for a statistical model (53). Upon selection, we ended up with the five ‘best’ models, representative of each of the four model options described above.

In a final step, we identified the ‘winning model’ among the 4 selected ones (based on the same criteria as the previous selection) and compared it with the following full model based on 6 regressors:

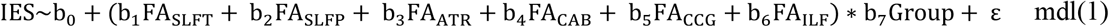

Where *Group* is a categorical variable describing the subjects’ group (ADHD vs TD).

We then tested how the winning model could account for ADHD symptoms (ADHD-RS), beta modulation (PI(β)) and modulation of drug related beta oscillatory response (ΔPI(β)).

## Results

While a total of 22 ADHD and 27 TD children underwent the MR session, only a smaller group of 18 ADHD and 26 TD successfully completed both MEG sessions (see Fig. 2 for the task). In the following section, we report the results of the latter group, where we quantified the association between white matter tracts microstructure, behavioral performance and MEG-derived measures. The bigger sample was considered for the association between diffusion weighted imaging (DWI) results and ADHD symptoms score.

### Behavioral performance

Behavioral performance in the ADHD group significantly improved following MPH administration (t_(17)_=2.49, *p*=.023), as reflected by lower inverse efficiency scores (IESs: accuracy/reaction time). No significant difference in IES was found between TD and ADHD: the TD group did not perform better when compared to the ADHD group in the Placebo (t_(42)_= −.22, *p*=.827) nor in the MPH condition (t_(42)_= .77, *p*=.445).

### Beta desynchronization in preparation to response to the cued target

As depicted in **Figure 6**, ADHD subjects in the placebo conditions exhibited lower overall beta depression, which was restored following MPH intake. This was observed as a diminished PI(β)s with values closer to those observed in the TD group (see Supplementary Methods and Materials). We will here focus on the association between beta modulation and white matter microstructural diffusion properties.

**Figure 6.**
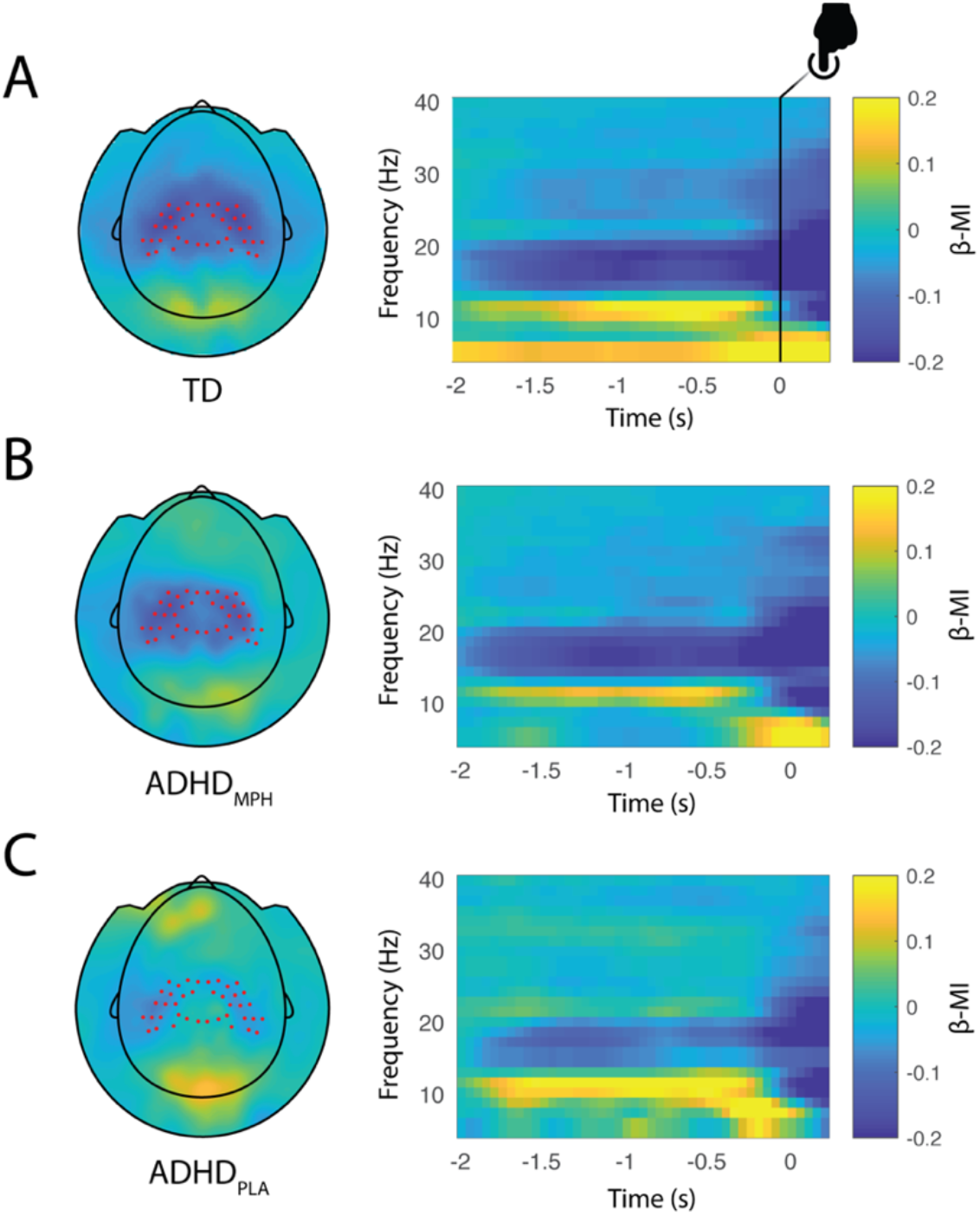
Beta modulation indices in the three conditions. (adapted from (1)),. Topographic plot (left) and respective time frequency representations (TFRs) (right panel) of power modulation (β-MI) for the typically developing group (TD) (A), ADHD_MPH_ group (B) and ADHD_PLA_ group (C). Red dots superimposed on the topographies denote sensors of interest as defined in Figure 4A. Notably, beta preparation is stronger in the TD group, while progressively decreases in the ADHD_MPH_ group, being weakest in the ADHD_PLA_ group.

### Definition of white matter structures of interest: model building strategy

Following the step-wise model selection criteria described in the *Materials and Methods* section, we identified a model with 3 regressors as the best fit (AIC= −67.34 and BIC=-53.09, Log Likelihood=42.67, Adjusted R^2^=.19) which included the regressors FA_SLFt_ (*p*=7 × 10^−4^), FA_SLFp_ (*p*=.003), and FA_ATR_ (*p*=.068). A 3D rendering of the three white matters tracts selected (SLFT, SLFP and ATR) is illustrated in **Figure 7** and they were considered as ROIs for later linear mixed model analyses.

**Figure 7.**
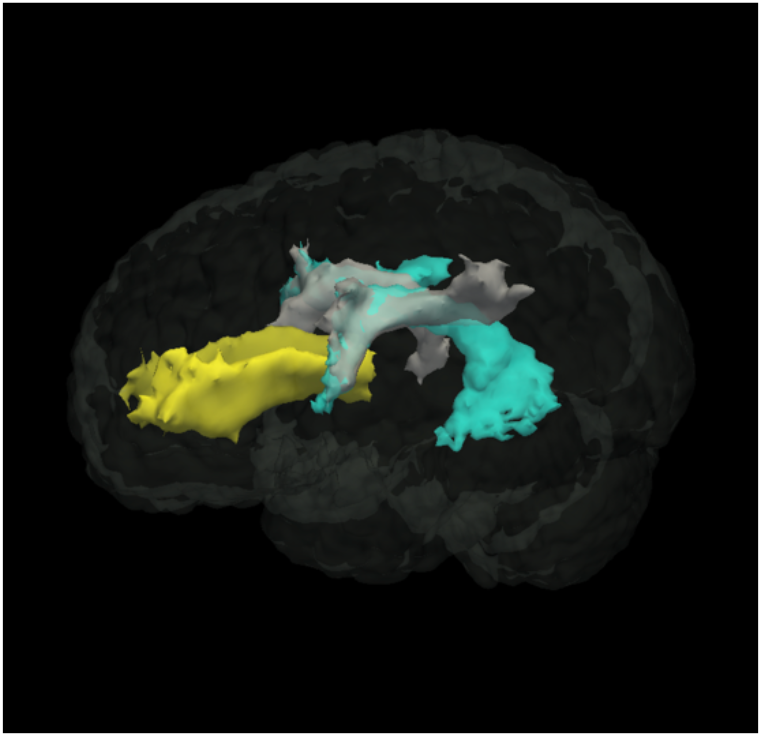
3D rendering of white matter ROI. Visualization of the probability distributions of the ROIs identified as tracts of interest according to the model selection approach: SLFp, SLFt and ATR.

We first enquired whether FA along these tracts differed between groups. We hence used a 2-way unbalanced ANOVA with factors *‘tract’* (ATR, SLFp, SLFt) and *‘group’* (‘TD’, ‘ADHD’) in relation to FA values. While the results yield a main effect of *group* (p=.009), showing a group difference in FA across all tracts, and *tract* (p=4.8× 10^−13^), reflecting a difference in FA between tracts, no significant interaction *group by tract* emerged (p=.453), indicating that the pattern of FA across tracts was similar across groups.

### Functional anisotropy of the Superior Longitudinal Fasciculus relates to behavioral performance

We considered task performance in relation to the three identified tracts of the winning model. Beta coefficients and adjusted response plots (i.e., showing the response as a function of one predictor, averaged over the others) derived from the model are shown in **Figure 8**: SLFt and SLFp were associated with beta coefficients of opposite sign: while higher FA values in the SLFt were associated with a higher IES (p= 7×10^−4^), i.e. worse performance (**Figure 8B**), a higher FA in the SLFp accounted for lower IES (p=.003), i.e. better behavioral performance (**Figure 8C**). A significant interaction term furthermore emerged for FA_SLFt_ and FA_SLFp_ with *Group* (p=.002 and p=.023, associated with a negative and positive coefficient, respectively). A negative interaction term of FA_SLFt_ with *Group* indicates that the effect of FA_SLFt_ on IES for the TD group was relatively weaker than for the ADHD group. *Vice versa*, a positive interaction term of FA_SLFp_ with *Group*, denotes that the effect of FA_SLFp_ for the TD group was relatively stronger than for ADHD.

**Figure 8–.**
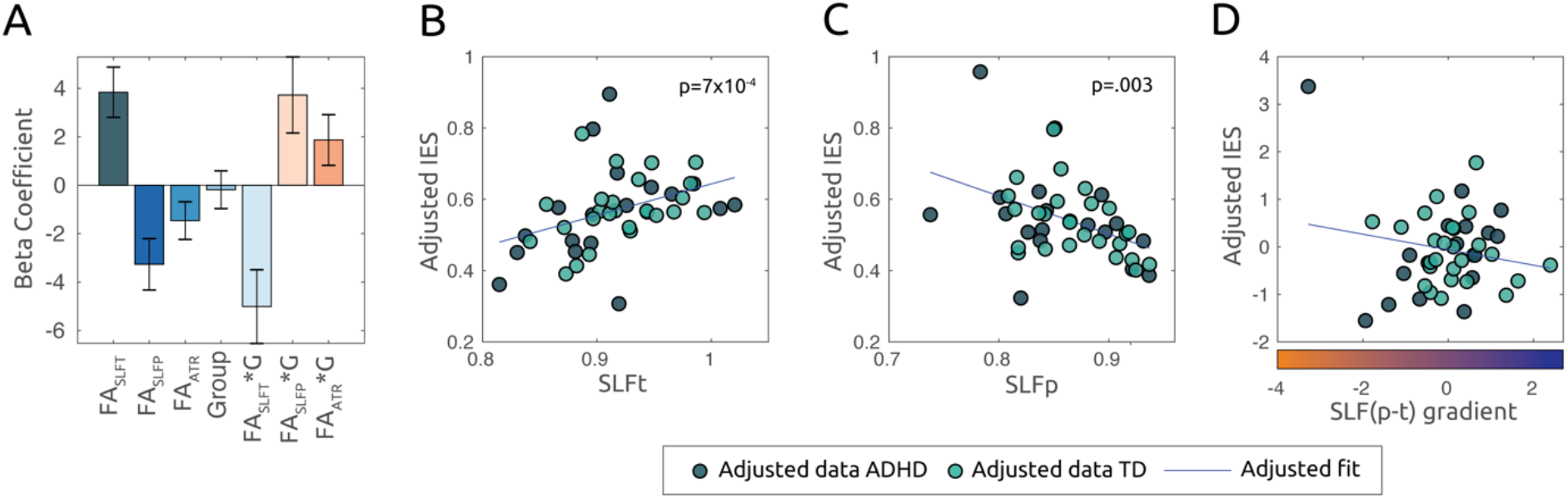
FA in SLFp and SLFt predict behavioral performance in the task. A. The bar plot displays the beta coefficients associated with the linear mixed model mdl(1), where mean FA values within the identified tracts of interest are set as explanatory variables for behavioral performance, as indexed by the IES. Error bars indicate standard error of the mean. The adjusted response plots in B and C show, respectively, the behavioral performance (IES) as a function of the FASLFt and, FASLFp, while averaging over other regressors in the model in (A). D. Adjusted response plot displaying the association between IES and parietotemporal gradient SLF(p-t), averaged over the residual regressors (mdl(1.a)). Positive SLF(p-t) values indicate higher FA along parietal as compared to temporal endings within the SLF.

Given the strong association between the two regressors FA_SLFt_ and FA_SLFp_ (r= .86, p=2×10^−4^; see correlation matrix in **Figure 5**) and based on the high degree of overlap between the two reconstructed tracts (as they overlapped anteriorly; **Figure 4,7**) more analysis was required to determine how SLFp and SLFt related to performance. We hence computed a measure describing the parietal-to-temporal SLF imbalance, according to:

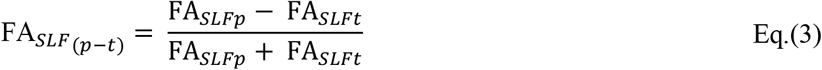

As a result, a given subject would display a specific degree of parietal-to-temporal diffusivity (i.e. imbalance) along the SLF tract: A higher value of the imbalance reflects a stronger diffusivity at parietal locations along the tract, and a lower value along the gradient reflected a stronger diffusivity at temporal locations.

We therefore considered a model specifically incorporating the parietal-temporal imbalance:

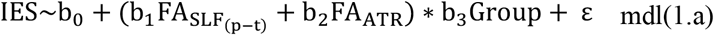

**Figure 8D** shows the association between IES and FA_SLF(p-t)_. As a corollary of Eq.(3), the negative partial association (p=.001) shows that subjects with a higher parietal than temporal FA along the SLF tract, were also the ones with a better behavioral performance. Also in this model, a significant interaction (p=.006) emerged between tract*Group, which reflected that the effect of _FASLF(p-t)_ on IES was stronger for the TD group compared to the ADHD.

### Fractional anisotropy of the ATR predicts ADHD symptoms

In order to assess whether the FA values of the white matter ROIs were related to symptoms (as measured by ADHD-Rating Scale; ADHD-RS), we considered the full sample of participants. We included symptoms across both the TD and the ADHD (placebo condition) group, hence embracing the notion that ADHD symptomatology derives from a ‘spectrum’, rather than from a dichotomous distinction with TD peers. The following model was then applied:

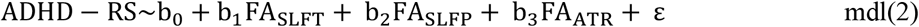

The resulting model was significant (R^2^=.22, p=.01, **Figure 9A**), and FA_ATR_ was associated with a significant partial coefficient (p=.007) showing a negative relationship with ADHD symptoms (**Figure 9B, 9C**): higher FA in the ATR predicted an overall lower ADHD symptomatology. FA_SLFp_ and FA_SLFt_ did not show a significant partial correlation with symptoms score (p=.494 and p=.309, respectively).

**Figure 9.**
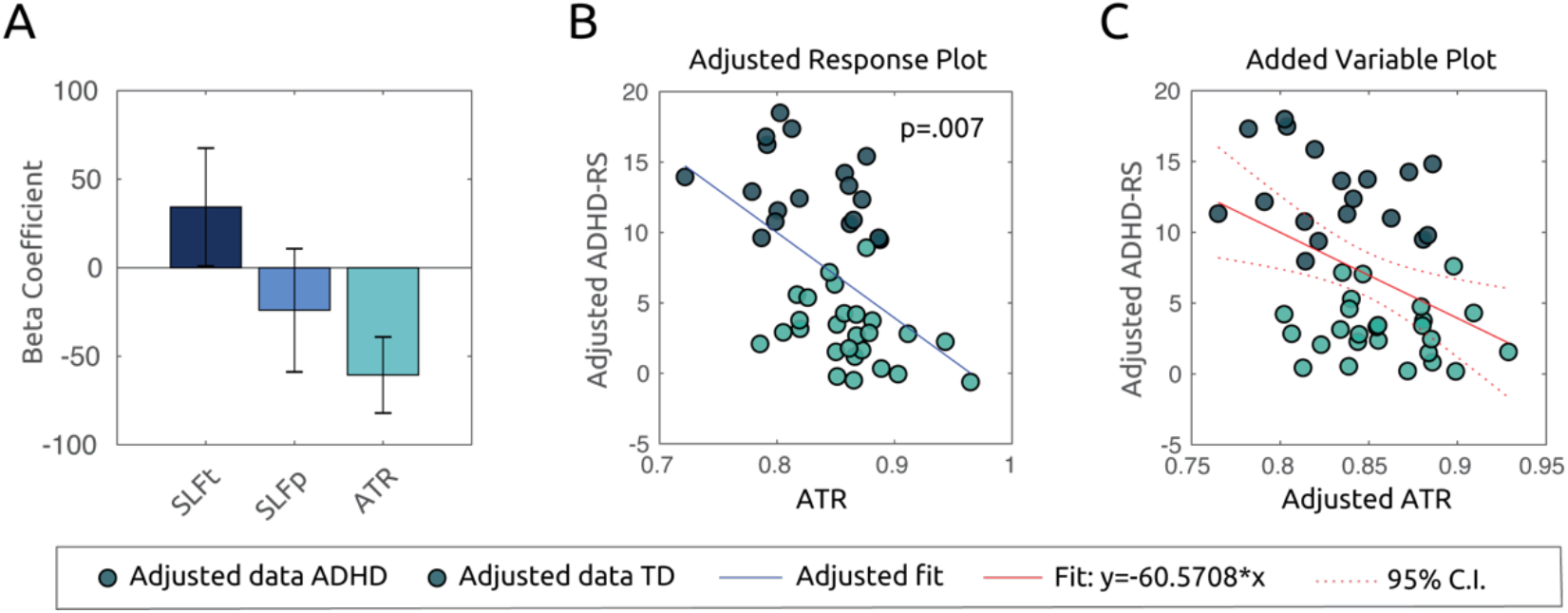
FA in ATR predicts ADHD symptoms severity in all subjects. A. Bar plot shows beta coefficients associated with mdl(2), where mean FA values along the tracts of interest are defined as predictors for ADHD-RS symptoms score in all subjects. FAATR is associated with a significant partial regression coefficient (p=.007). Scatter plots in B and C show, respectively, adjusted response plot averaged over the other predictors and the added variable plot (partial regression), of symptoms as a function of FAATR. Lower diffusivity along the ATR corresponded to higher average ADHD symptomatology along the spectrum.

### Parietotemporal gradient along the SLF reflects methylphenidate’s effect on preparatory beta depression

Finally, we sought to investigate the relationship between FA values in the selected ROIs, and the patterns of MPH associated beta depression (PI(β)). As described in (1), MPH intake normalized aberrant beta depression in ADHD, initially lower as compared to controls. In a first model, we aimed at identifying whether a combination of FA values in the selected ROIs accounted for the degree to which subjects were able to suppress their somatosensory beta power in preparation to a motor response (PI(β)). To this end, we considered PI(β)s for TD subjects and ADHD subjects in the placebo condition as dependent variable in the following model:

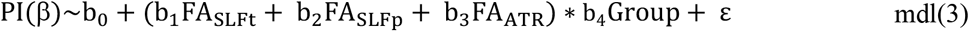

The resulting overall model was not significant (p=.762), neither were the main or the interaction effects in the regressors (FA_SLFt_ : p=.890; FA_SLFp_ : p=.985; FA_ATR_ : p=.754, FA_SLFt_*Group : p=.934, FA_SLFp_*Group: p=.689, FA_ATR_*Group : p=.655).

Next, given the effects of MPH on modulations of the beta oscillations within the ADHD group, we enquired whether, instead, a linear combination of the ROIs’ FA properties, could explain the *changes* in beta depression following medication intake (ΔPI(β)). We hence implemented the following model, consistently with the principles abovementioned:

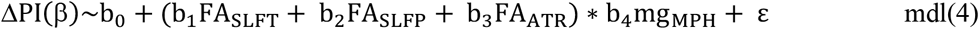

Here, we added the term mg_MPH_ as a categorical variable to control for the dosage of MPH administered prior to the task (15/10mg). The resulting model was significant with R^2^=.83 and p=.003 (**Figure 10A**). Analyses of main effects showed a significant negative association between FA_SLFt_ and ΔPI(β) (*b*_1_: p=.001) (**Figure 10B**), and a positive association between FA_SLFp_ and ΔPI(β) (*b*_2_: p=1×10^−4^) (**Figure 10C**), while no main effects for FA_ATR_ and mg_MPH_ were found (p=.910 and p=.693, respectively).

**Figure 10.**
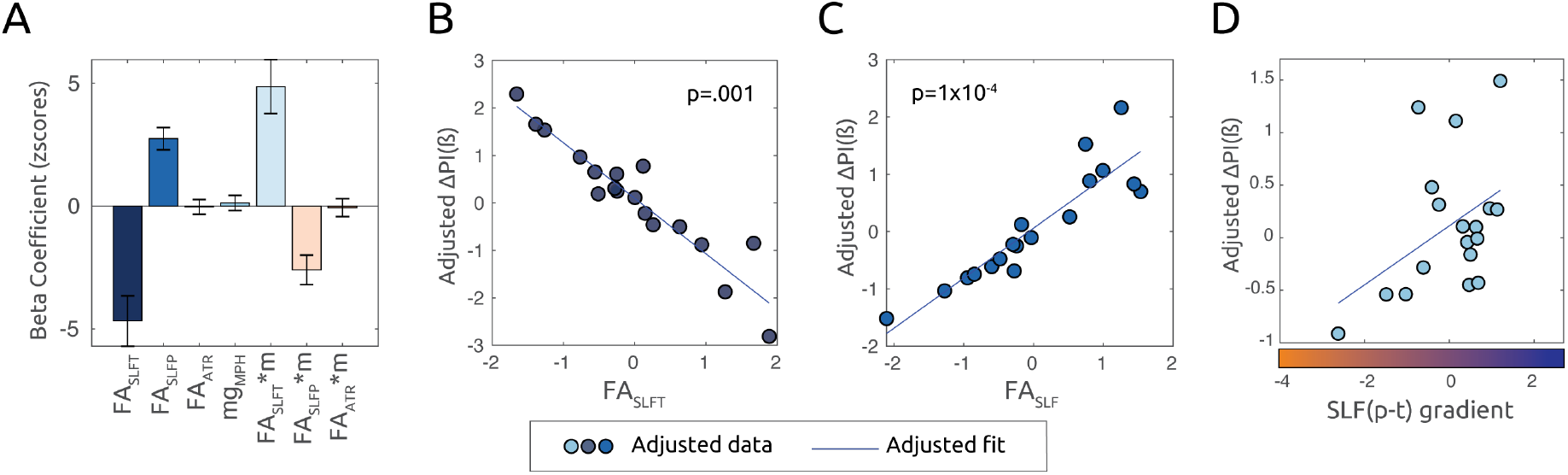
FA in SLFp and SLFt predict MPH effects on β depression in the ADHD group. A. Bar plot shows beta regression coefficients associated with mdl(4), where mean FA values along the tracts of interest are defined as predictors for changes in beta modulation due to medication intake (ΔPI(β)). According to Eq.(2), a higher ΔPI(β) value for a given ADHD subject reflects bigger changes in depression of beta oscillations following medication intake. B. Adjusted response plot showing ΔPI(β) as a function of FASLFt. A significant beta coefficient (p=.001) reveals that FASLFt predicted lower changes in β power depression due to MPH. C. Adjusted response plot showing ΔPI(β) as a function of FASLFp. Higher FASLFp corresponded to stronger β depression changes following MPH intake (p=1×10-4). D. Adjusted response plot illustrating ΔPI(β) as a function of parieto-temporal SLF gradient according to mdl(4.a). Higher SLF(p-t), for a given ADHD subject, reflected higher diffusivity at parietal endings, as compared to temporal endings, of the SLF. The significant positive relationship (p=7×10-4) suggests that the gradient of parieto-temporal diffusivity in the SLF is predictive of the effects of MPH on β depression.

The main effects in the model denoted a bigger MPH effect for subjects displaying higher FA in the SLFp, while subjects displaying higher FA in the SLFt were the ones whose sensorimotor beta was less affected by MPH administration.

Although the model produced significant interaction terms between FA_SLFt_ and FA_SLFp_ with mg_MPH_ (p=.001 and p=.002, respectively), we did not pursue any post-hoc examination of such effect, given a very low ratio between 10/15mg dosage in the sample (0.38) would not produce statistically reliable results.

According to the same principle which led to mdl(1.a), we merged the two regressors FA_SLFp_ and FA_SLFt_ into the gradient denoted as FA_SLF(p-t)_, and considered an equivalent to mdl(4) as follows:

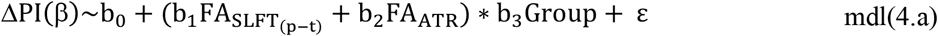

In **Figure 10D** we present an alternative and equivalent representation of the plot in **Figure 10B and C**, where the linear association between ΔPI(β) and FA_SLF(p-t)_ is presented. Here, we can observe a significant positive partial association (p=7×10^−4^) which indicates that subjects with a higher parietal compared to temporal FA along the SLF tract, were also the ones whose beta depression was more affected (enhanced) by MPH.

## Discussion

In the current study, We employed DTI to estimate microstructural properties of main bilateral fasciculi and explored their role in mediating behavior and beta power modulation associated with ADHD symptomatology.

We first showed that, in both groups, lower values of fractional anisotropy within the ATR were related to ADHD symptom severity and that parieto-to-temporal FA-imbalance within the SLF accounted for behavioral performance in the attentional task. Importantly, in the ADHD group, the same SLF gradient was predictive of the effects of MPH on beta power modulation.

The ATR originates from anterior and medial nuclei of the thalamus and radiates along the anterior thalamic peduncle and the anterior limb of the internal capsule to reach the frontal cortex (54,55). The thalamocortical feedback loop is crucial in conscious processing, and its role in attention is to provide a functional link between otherwise structurally segregated cortical areas, supporting different aspects of attentional selection and working memory (56,57). Prior morphological analyses have shown that ADHD is associated with altered shape and volume of thalamic nuclei, whose connections within several cortical regions, particularly frontal areas, seem disrupted in affected children (58,59). On the other hand, there is still controversy over the validity of thalamic volume as anatomical correlate of the disorder, given that case-control volumetric differences in this area were not confirmed in a recent mega-analysis (60). However, that mega-analysis did provide evidence for a different influence of age on thalamic volumes between clinical and non-clinical samples. The thalamus is a rather complex structure composed of a set of cytoarchitectonically segregated nuclei, each providing specific thalamocortical signals and relying on partially independent neural circuitry to mediate different cognitive functions. Hence, a more valid investigation of thalamic involvement in the pathophysiology of ADHD should take into account an specific analysis of such diverse nuclei (61), so far still absent in the ADHD literature. Arguably, currently the most convincing effects with regard to thalamus role in ADHD, are found in the context of connectivity studies, pointing to the importance of fronto-striatal circuits and their disruption in association to the symptomatology (62). Here, we corroborated these findings by embracing a different approach: the association between thalamocortical diffusivity and symptom severity we report, exploits the inherent heterogeneity in FA values among subjects, by linking this variance to behavioral symptomatology along the broader clinical spectrum. In other words, our results speak to the importance of thalamocortical connections in the interaction between attention and premotor functions which, if anomalous, are at the basis of clinical behavioral symptoms (as observed in ADHD).

We postulate that the strong interconnections between the thalamus and striatal regions, among which different basal ganglia nuclei, is one of the core variables into consideration when aiming at identifying structural correlates of ADHD. Reinforcing this finding, the TD group in our study displayed stronger anisotropic diffusivity along the ATR as compared to the ADHD group. Such notion further highlights the role of the basal ganglia and their morphological and volumetric differences as potential predictors of the disorder (60,63–68).

The second main and novel finding related the parieto-to-temporal FA-imbalance within the SLF with behavioral performance in the attentional task (in both groups): a higher FA in the parietal-SLF compared to the temporal-SLF predicted faster and more accurate responses in both ADHD and controls. Furthermore, in the ADHD sample, the same gradient explained the effects of MPH on beta modulation: individuals displaying higher FA in the parietal than the temporal SLF were also the ones whose beta power during response preparation increased more with MPH. FA along the SLF likely reflects the functions of the frontoparietal control network (FPCN). This network is one of the core anatomical components providing the basis of flexible attentional adaptations to different task demands (69). While the above results linking FA imbalance within the SLF to behavior and beta oscillations are seemingly orthogonal to the previous finding relating FA_ATR_ to symptoms severity, the former are not independent from thalamo-frontal influences. Indeed, the dopaminergic regulation of the prefrontal cortex and the striatum has been proposed to mediate the interaction between the different attentional systems (27,70,71), some of which structurally rely on the SLF.

Dopaminergic availability is modulated by MPH (21,24,72), which blocks the reuptake of the neurotransmitter, hence allegedly increasing the functional interactions within attentional networks. Given these premises, it is not surprising that diffusivity along the superior longitudinal fasciculus favors the effects of MPH: a stronger connectivity, as indexed by anisotropy along the tract, promotes communication between frontoparietal areas, which is further maximized by the stimulant’s action. Crucially, we found that an imbalance of FA in favor of parietal rather than temporal regions is associated with stronger effects of medication on beta modulation. Important nodes of the FPCN are found in dorsolateral prefrontal cortex, frontal eye fields and intraparietal sulcus (73), regions that are connected by the dorsal fibers of the SLF (74). Previous studies have already proposed that the mechanisms of action of stimulant medication in ADHD are strongly reflected by its activity on frontoparietal regions (18,75), whose under-activation is one of the neural correlates of the disorder (76–78).

It is important to consider the link between dopaminergic coordination of attentional networks and increased beta modulation. Evidence from Parkinson’s Disease (PD) patients has provided a theoretical framework according to which dopamine levels within the basal ganglia-cortical loop have a direct influence on beta oscillations (79,80). Anomalous beta oscillations are highly correlated with PD pathology (81), with evidence from in vivo recordings and mathematical models suggesting they originate from dopamine degeneration in cortical projections to the striatum (82–84). Coupled with findings of increased striatal dopamine transporter (thus lower dopaminergic availability) in ADHD (85–88), we here propose an attentional network which relies on the dopaminergic input from thalamo-striatal regions to the prefrontal cortex, which in turn mediates the activity along the FPCN. Higher FA along frontoparietal tracts tends to reflect increased connectivity and is being investigated as indirect tool to infer dopaminergic functions in the pathological brain (89–91). Relatively stronger parietal FA may thus underly a facilitatory mechanisms for MPH action in the brain, as revealed by a stronger increase in preparatory beta desynchronization in children displaying higher parietal-to-temporal FA gradient.

Although limited in its sample size, this study offers important new insights in the potential of multimodal imaging to investigate and identify the sources of attention performance in the brain: by coupling evidence from electrophysiological measures and information about white matter integrity, we are able to investigate the origin of aberrant brain activity observed in ADHD and improve our understanding of the mechanisms of action of stimulant medication. Furthermore, it might encourage future research on the issue, clarifying whether FA represents a predictor of MPH responsiveness or it is rather one of the neural targets of medication, as could potentially be inferred by longitudinal research.

## Disclosures

Jan K Buitelaar has been in the past 3 years a consultant to / member of advisory board of / and/or speaker for Janssen Cilag BV, Eli Lilly, Medice, Takeda/Shire, Roche, and Servier. He is not an employee of any of these companies, and not a stock shareholder of any of these companies. He has no other financial or material support, including expert testimony, patents, royalties. The remaining authors declare no competing and financial interests.

## Acknowledgements

The authors gratefully acknowledge the support of the Marie-Curie ITN grant ChildBrain (grant number 641652).

## Notes

### Competing Interest Statement

The authors have declared no competing interest.

