## Supplementary Material for "Dorsal-to-ventral imbalance in the superior longitudinal fasciculus mediates methylphenidate’s effect on beta oscillations in ADHD"

#### Supplementary Methods and Materials

***Participants***

ADHD children were recruited from different local establishments and general practitioners' offices. Children from both groups were recruited via advertisements in local schools and public places, and through the study website. The experiment was conducted in compliance with the declaration of Helsinki and was approved by the local ethics board (CMO region Arnhem-Nijmegen, 2016-2268) and study was preregistered on the Netherlands Trial Register (NL56007.091.15). All parents gave written informed consent, while children gave oral consent.

For both groups, inclusion criteria were: 1) age between 8 and 12 years at the time of the experiment; 2) male; 3) enrolled in primary, not secondary, school; and 4) derived FSIQ measure > 70. For the ADHD diagnosed group, additional inclusion criteria were: 1) a clinical diagnosis of ADHD according to DSM-5 criteria (American Psychiatric Association, 2013); 2) scoring in the clinical range on the ADHD DSM-5 rating scale; 3) being treated with stimulant medication for ADHD (either long or short active formulations), which started at least three months before the inclusion in the study. For both groups, exclusion criteria were: 1) neurological disorders (e.g., epilepsy), currently or in the past; 2) cardiovascular disease, currently or in the past; 3) serious motor or perceptual handicap; 4) standard MRI exclusion criteria (e.g., metal objects/fragments in the body, active implants, brain surgery, claustrophobia). Comorbidities were documented, where present, as assessed by the Childhood Behaviour Checklist (CBCL) (Achenbach, 1999), completed by the parents. For the ADHD group, additional comorbidities screening was performed during the psychiatric intake. For the TD group, the absence of psychiatric disorders was assessed via a screening form completed and signed by parents. 27 children with a diagnosis of ADHD and 27 typically developing (TD) male children were included in the current study. 9 children in the ADHD group withdrew from the experiment after at least one session, because of the parents' decision (N=5), claustrophobic reaction to the MRI scan (N=1), claustrophobic reaction in the MEG room (N=1) and excessive moments during MEG measurements (N=2). One participant in the TD group was excluded from the analysis, given he was diagnosed with ADHD following the end of the data collection.

Continuous MEG data were acquired and analyzed for the remaining 44 participants, leaving a total of 18 children in the ADHD group (age: 10.8±1.0 years) and 26 children in the TD group (age: 10.7±1.2). Although structural MRI data were collected for 50 children (27 TD), we here focus only on electromagnetic results. The MRI protocol and data analysis are thus not further described.

While only the completed datasets (MEG and MRI/DWI) were considered for the analyses related to behavioral task performance and MEG-recorded beta modulation (leaving a total of 18 children diagnosed with ADHD (mean age 10.8±1.0 years) and 26 TD children (10.7±1.2)), all MR data (49, of which 27 TD: mean age 10.7±1.3, and 22 ADHD mean age 10.7±1.0)) available were instead used for analysis of correlation with ADHD symptom scores.

***Experimental Design***

The study consisted of two experimental sessions for the TD group, and three sessions for the ADHD group, within a randomized placebo-controlled double-blind crossover design. Before visiting the institute, parents were asked to fill the Child Behavioral Checklist (CBCL) and ADHD rating scale questionnaires (ADHD-RS). Parents of the participants in the ADHD group, were asked to fill the latter one twice: once referring to the child's symptomatology pattern while observing him on medication, and once referring to behavior without medication (in this case, the parents were asked to observe their child during the withdrawal period before the experiment). During the first intake-session, children of both groups underwent behavioral testing. This involved a *Line Bisection Test* (LBT), to assess individuals' spatial biases by asking participants to mark with a pencil the center of a series of horizontal lines (performed both with the left and the right hand) and, if intelligence had not been assessed over the past two years, the *Vocabulary* and *Block Design* subtests of the Wechsler Intelligence Scale (WISC-III), designed for children (Kaufman, 1994; Woolger, 2001); Dutch version in (Kort et al., 2002)), in order to estimate the FSIQ. These subscales have been shown to hold high correlation with the full-scale IQ testing (Herrera-Graf et al., 1996). For the ADHD group, an in-depth intake was conducted by a psychiatrist, where a 30 min interview with parents and son, together with physical examination, were employed to determine the medication dosage to be used during the task, based on operating procedures followed in prior studies, describing a medium dosage of 0.3 mg/kg (Linssen et al., 2014). Based on the screening, one of the two standardized dosages was chosen (either 10 or 15mg Methylphenidate immediate release; IR-MPH). Following the behavioral screenings, children of both groups were introduced to the MRI procedure, and were given the opportunity of getting acquainted with the scanner environment by practicing in a so-called '*dummy scanner*', which simulates the environment of a real MRI scanner. If the child was unable to stay still or felt uncomfortable during the dummy scan, the session was cancelled and he was excluded from further participation. Otherwise, the MRI session took place for a duration of ~30mins.

During the second visit, both groups undertook the MEG testing, followed by 3-D head digitization using an electromagnetic digitizer (*Polhemus*, Colchester, VT). For the TD group, this constituted the last day of testing, while for the ADHD group, two MEG sessions were planned at two different visits, separated by at least one-week interval. Children diagnosed with ADHD performed the MEG task twice under two conditions (MPH and placebo), according to a randomized order and double-blind procedure. Prior to each MEG session, participants were asked to withdraw from their clinically prescribed medication intake for 24 hours, depending on the planned time of the recording session (e.g. if the session was planned in the morning of Monday, the child took his last medication on Sunday morning, abstaining until the experiment). To ensure this, parents agreed on being contacted prior to the experiment to be reminded about the medication withdrawal procedure, to be followed in preparation to the testing day (e.g., in the example above, parents were contacted on Saturday). The 24 hour treatment suspension allowed to control for withdrawal symptoms related to drug administration (rebound effect) (Carlson and Kelly, 2003). MEG testing began one hour after medication intake, allowing to reach on average moderate plasma concentration ($C_{max}$) of the drug along the experiment, which progressively increases and reaches its peak around the second hour post-intake (Quinn et al., 2007). After completion of the MEG session, participants were asked to proceed with their regular treatment using their own stimulant formulation. At the beginning of the experimental session, parents and children were asked to confirm that the medication withdrawal procedure was followed appropriately.

***MEG data acquisition and analysis***

Electromagnetic brain activity was recorded from the participants seated in a CTF 275-sensor whole-head MEG system with axial gradiometers (CTF MEG Systems, VSM MedTech Ltd.). The data were sampled at 1200Hz, following an antialiasing lowpass filter set at 300Hz. Head position was constantly monitored throughout the experiment via online head-localization software. This was done by three head localization coils placed at anatomical fiducials (nasion, left and right ear), allowing, if necessary, readjustment of the participant's position between blocks.

MEG data analysis was performed using the MATLAB FieldTrip Toolbox (Oostenveld et al., 2011). The continuous data were segmented in epochs time-locked to the onset of the motor response (-2000 to 200 ms). A notch filter was applied at 50, 100, 150 Hz to remove line noise, after which the mean was subtracted and the linear trend removed. Trials with incorrect responses according to the cued hemifield were discarded. Artifacts were rejected first via a semi-automatic artifacts rejection of trials with MEG sensor jumps and muscle artifacts, then through visual inspection to further detect and remove trials with eye blinks and systematic saccades which did not exceed the boundary box set by the eye tracker but still produced detectable visual artifacts. ICA was then used to further remove components reflecting eye blinks and heart artifacts.

Prior to the time-frequency analysis of power, we generated virtual planar gradiometers from spatial derivatives of the axial magnetic components (Bastiaansen and Knösche, 2000). Time-frequency representations (TFRs) of power were then computed for each pair of orthogonal planar gradiometers, and power values were then summed for each MEG sensor. Power analysis was performed using a 600 ms time-window sliding in steps of 50 ms along the time window of interest of 2200ms length specified above, locked to the onset of the motor response. The resulting data segments were multiplied by a Hanning taper and a fast Fourier transform was applied in the 2 – 40Hz frequency range, in steps of 1.66 Hz. The steps above were also followed for cue-locked epochs, obtained by re-defining the trials according to the onset of the cue, considering the time window of the preceding 1000ms and the following 1000ms.

In order to estimate patterns of beta modulation indices (β-MI) in preparation to the motor response, we performed a baseline correction (relative change) of response-locked data segments with respect to pre-cue activity per sensor. For each subject, we applied a relative baseline correction with respect to the averaged power in the time window [-300 – 0] ms prior to the cue. The baselined TFRs for all subjects were then averaged across conditions and the three groups (TD, $ADHD_{MPH}$, $ADHD_{Placebo}$), in order to identify the sensors of interest to be used in further analyses. To this aim, we selected a cluster of 20 symmetrical pairs of central sensors, displaying the highest beta depression values (lowest β-MI) in the 1000ms time-window preceding the onset of the motor response. The same sensors were then used for the estimation of mean beta desynchronization indices (Preparation Index of beta, PI(β)) for each subject, by averaging MI(β)s across the time window of interest ($f$= 15 – 30Hz, -1000 < t < 0 ms).

***MRI data acquisition parameters***

MRI data were acquired at the Donders Institute (DCCN) using a 3T MAGNETOM Skyra MR scanner (Siemens AG, Healthcare Sector, Erlangen, Germany) with a product 32-channel head coil. The MRI protocol included a T1-weighted MRI scan for anatomical reference and analysis and diffusion-weighted MRI scans for probing microstructural properties and for performing fiber tractography.

Whole brain high resolution T1-weighted anatomical data were acquired with sagittal as primary slice direction (MP-RAGE, 192 slices, acquisition matrix 256×256, voxel size 1×1×1mm, slice thickness 7.0 mm, TR=2300ms, TE=3.03ms, TI=1100ms, flip angle= 8°, GRAPPA-acceleration 2). Whole brain diffusion-weighted images (DWI) were collected with the following protocol: multiband factor 3, reverse phase-encoding polarity, acquisition matrix: 104x104x72, 111 diffusion-weighted directions; b-factor 1500s/mm2; 11 non-diffusion- weighted images; interleaved slice acquisition; TE/TR=99.8/3670ms; multi-band acceleration 3; voxel size 2×2×2mm, EPI factor=104, Echo spacing=0.73ms). The approximate total recording time for the MR session was 27 minutes.

### Supplementary Discussion

#### *Study Limitations*

Given the study design, our findings do not allow to disambiguate the directionality of cause-and-effect: white matter changes could occur in response to stimulant treatment, resulting in the association between higher FA along the SLF and stronger MPH response. Alternatively, the higher diffusivity along this tract may boost the effects of stimulants, as reflected by higher changes in beta power.

Furthermore, in vivo DTI tractography as the one implemented in the context of the current paper, provides a fiber tracts model, which does not directly reproduce physiological fiber connections. As previously stated, we implicitly rely on the assumption that FA reflects the underlying tissue microstructure, such as integrity of myelin sheath, which impacts the overall mobility of water along axons (33).

A potential limitation can be identified in the sample size. While a modest number of participants was achieved in the TD group, close to the one initially planned for the study, practical complications and a higher burden impacted the ADHD group, which, based on the study design, undertook more hours of testing. Further time and ethical constraints did not allow for a prolongation of the intake process in order to reach a bigger sample in the ADHD group.

From a statistical standpoint, the model selection strategy we applied partially deals with the problem, by parsimoniously providing a trade-off between overfitting and underfitting. Our implementation is still liable to overfitting due to the fact that model testing is applied to the same sample, although with different dependent variables. Our findings will likely benefit from an investigation in an independent sample, which could further corroborate the associations between structural and electrophysiological signatures of attention that we presented.

Importantly, a caveat applies to the generalization of our current results and consequent clinical implications, which cannot extend to adult populations and females until validation in the respective representative samples. On the other hand, stricter inclusion criteria and control over the heterogeneity among study participants is often a strength when aiming at identifying neural biomarkers of a disorder.
